# Advantages of acute brain slices prepared at physiological temperature in characterization of synaptic functions

**DOI:** 10.1101/845461

**Authors:** Kohgaku Eguchi, Philipp Velicky, Elena Hollergschwandtner, Makoto Itakura, Yugo Fukazawa, Johann Georg Danzl, Ryuichi Shigemoto

## Abstract

Acute brain slice preparation is a powerful experimental model for investigating the characteristics of synaptic function in the brain. Although brain tissue is usually cut at ice-cold temperature (CT) to facilitate slicing and avoid neuronal damage, exposure to CT causes molecular and architectural changes of synapses. To address these issues, we investigated ultrastructural and electrophysiological features of synapses in mouse acute cerebellar slices prepared at ice-cold and physiological temperature (PT). In the slices prepared at CT, we found significant spine loss and reconstruction, synaptic vesicle rearrangement and decrease in synaptic proteins, all of which were not detected in slices prepared at PT. Consistent with these structural findings, slices prepared at PT showed higher release probability and higher detectability of long-term depression after motor learning compared with slices prepared at CT. These results indicate substantial advantages of the slice preparation at PT for investigating synaptic functions in different physiological conditions.

## Introduction

The living acute brain slice preparation has been developed and extensively used as a powerful experimental model for investigating the structural and functional characteristics of synaptic connectivity of neuronal circuits in the brain^1–5^. The acute slice preparation is readily accessible for electrophysiological and optical recording, and expected to retain the cytoarchitecture and synaptic circuits *in vivo* except for the long-range projections. In general, to prepare acute brain slices, a whole brain is dissected out from an animal and quickly immersed into ice-cold (< 4 °C) cutting solution to slow down the metabolic activity in tissue blocks, which are sliced by a microtome at ice-cold temperature (CT). The slices are then pre-incubated in artificial cerebrospinal fluid (ACSF) warmed at physiological temperature (PT, 35-37 °C) for up to 1 hour to recover the neuronal activities prior to electrophysiological or optic recordings^5,6^. The preparation method of acute brain slices at CT, however, causes alterations of molecular and cellular components of neurons. For example, exposure of hippocampal slices to CT induces disassembly of microtubules and eliminates dendritic spines in neurons^7,8^. Although re-warming of the hippocampal slices revives microtubule structures in neurons, excessive proliferation of the dendritic spines results in a higher density of synapses than that observed before cooling. Slicing at CT and subsequent incubation at 37 °C also reduces the protein level of α-amino-3-hydroxy-5-methyl-4-isoxazolepropionic acid-type glutamate receptors (AMPARs), which mediate synaptic transmission and plasticity, in lysates of the acute hippocampal slices of rats^9^. These artificial modifications may cause disadvantages of brain slicing at CT for investigating synaptic features.

Recently, a method for acute slice preparation at PT has been developed to improve the quality of cerebellar slices in aged rodents^10^. In the cerebellar slices from aged rat (> 2 months-old) prepared at PT, Purkinje cells (PCs) survived better than those in slices prepared at CT without altered intrinsic excitability of the cells. Another study reported that the slice preparation at PT enhanced the cellular viability and mitochondrial activities in rat hindbrain slices compared with the preparation at CT^11^. Although these reports suggest some advantages of the acute slice preparation at PT (warm-cutting method), quantitative differences in electrophysiological, molecular biological and ultrastructural properties of synapses in the brain slices prepared at CT and PT have not been examined.

To clarify which parameters of synaptic architecture and functions are affected in slices prepared at CT and PT, we investigated various ultrastructural and electrophysiological features of synapses in acute cerebellar slices prepared at CT and PT using electron microscopic (EM), super-resolution microscopic and electrophysiological techniques in acute cerebellar slices. We show that the conventional cold-cutting method causes loss and reemergence of dendritic spines, rearrangement of synaptic vesicles in nerve terminals and decrease in both pre- and postsynaptic proteins such as AMPARs and P/Q-type voltage-gated Ca^2+^ channels (Cav2.1). In contrast, the acute cerebellar slices prepared at PT showed no significant changes in these features during the recovery time, and also less difference from perfusion-fixed tissue, indicating that the warm-cutting method can preserve the synaptic functions through the preparation process better than the cold-cutting method. We also demonstrate that long-term depression (LTD) in mouse cerebellum caused by adaptation of horizontal optokinetic response (HOKR) is better preserved in slices prepared at PT than at CT,indicating that the synaptic plasticity induced *in vivo* may be partially reset by the cold-cutting method through the loss and reconstruction of spine synapses. Altogether, our quantitative analysis suggests strong advantages of the warm-cutting method over the conventional cold-cutting method for investigating synaptic functions. This method facilitates a wide range of neuroscience research, especially for synaptic plasticity induced *in vivo.*

## Results

### Influence of the temperature during slice preparation on electrophysiological properties in synaptic transmission

We first examined whether different temperatures during slicing affect electrophysiological properties of synaptic transmission in parallel fiber-Purkinje cell (PF-PC) synapses in mouse cerebellum using whole-cell patch-clamp recording. The cerebellar slices (coronal, 250-300 μm thick) were prepared in ice-cold (< 4 °C) or warmed (35-37 °C) sucrose-based cutting solution to minimize the disruption of the spine structure^8^, and then pre-incubated in normal ASCF warmed at 37 °C for 1 h as a recovery time. Electrical stimulation applied to the PF in the molecular layer evoked an excitatory postsynaptic current (evoked EPSC) on the recorded PC. The amplitude of the evoked EPSCs increased depending on the stimulation intensity showing no significant difference between the two temperatures (Fig. 1a-b, Table.S1). The mean amplitude, 10-90% rise time, or decay time constant of evoked EPSCs also remained unaffected (Fig. 1c, Table.S1).

**Fig. 1.**
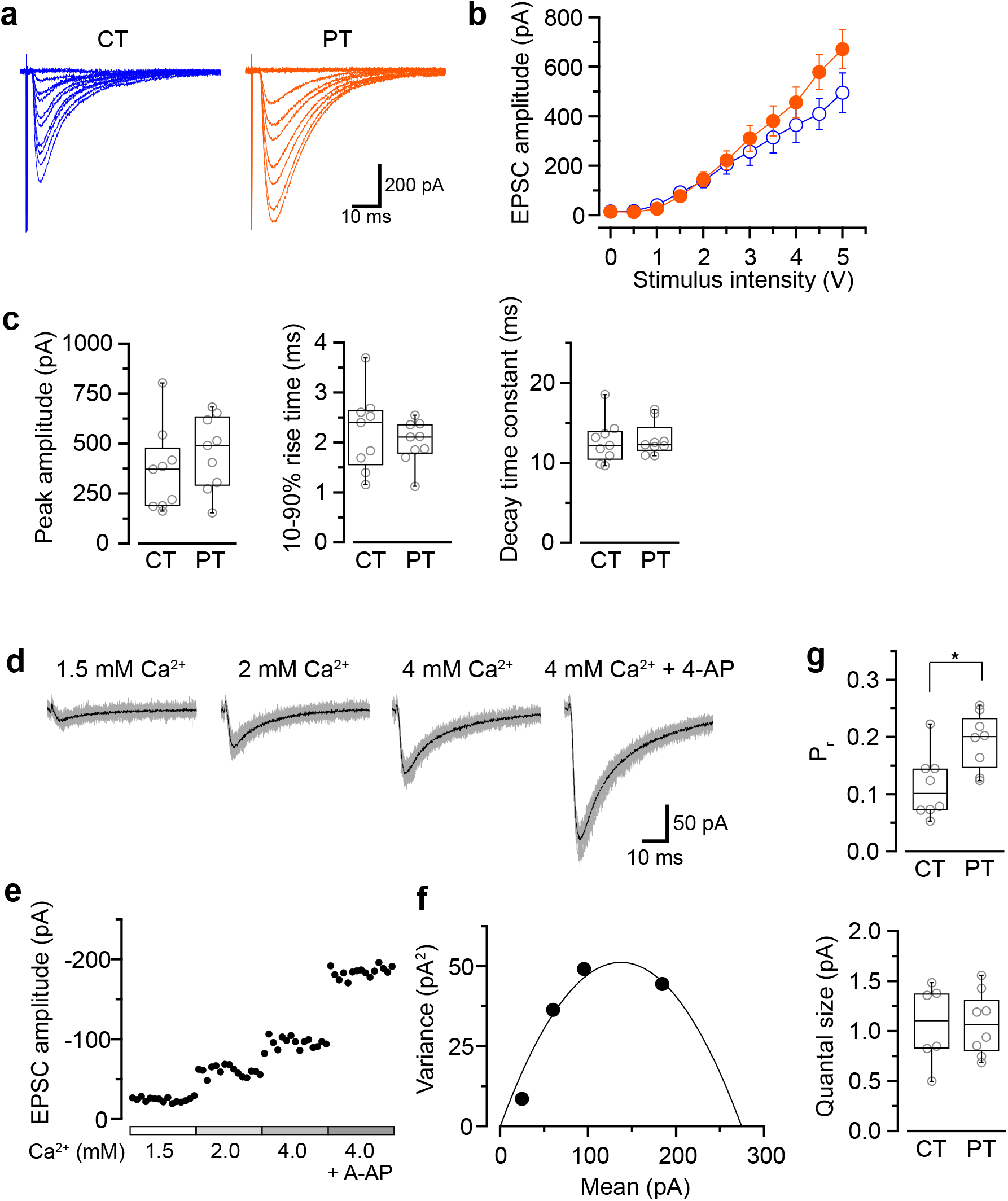
Influence of the temperatures during slicing on synaptic transmission in PC-PF synapses. **a**, Representative evoked EPSC traces elicited by several stimulus intensities (0-5 V, 0.5 V steps, superimposed) recorded from PCs in the slices prepared at CT (left) and PT (right). **b**, The relationship of evoked EPSC amplitude and stimulus intensity recorded from PCs in the slices prepared at CT (blue, n = 9 cells) and PT (orange, n = 9 cells). **c**, Summary of the peak amplitude (left), 10-90% rise time (middle) and decay time constant (right) of EPSCs (evoked by a 4-V stimulation) in cold- and warm-cut cerebellar slices. Each scatter indicates the mean value obtained from an individual PC (n = 9 cells for each). No significant difference was detected between CT and PT (amplitude: P = 0.34, rise time: P = 0.49, decay time constant: P = 0.82, n = 9 cells each, Welch’s *t*-test). **d-e**, Representative evoked EPSC traces (**d**) and plots of the amplitude (**e**) in various [Ca^2+^]_out_ recorded from a PC in the slice prepared at PT. Gray traces in (**d**) show the individual 15 traces and black traces indicate their averages. **f**, Mean-variance (M-V) relationship of evoked EPSCs obtained from the amplitudes shown in (**e**). The plot was fitted by a parabola equation. **g**, Summary of the release probability at 2 mM [Ca^2+^]_out_ (P_r_, top) and the quantal size (bottom) of PF-PC synapses in slices prepared at CT or PT estimated by M-V analysis. Each scatter indicates the mean value obtained from an individual PC. P_r_ recorded from the warm-cut slices (n = 8 cells) shows a significantly higher value than that from the cold-cut slices (n = 8 cells, P < 0.05, Welch’s *t*-test), whereas no significant difference in quantal size was detected between slices prepared at CT and PT (CT: n = 6 cells, PT: n = 8 cells, P = 0.96).

We then compared release probability (P_r_) of PF-PC synapses between the cold- and warm-cut slices using mean-variance (M-V) analysis^12,13^. For the parabola fitting, P_r_ was altered by simply changing extracellular Ca^2+^ concentration ([Ca^2+^]_out_, 1.5 to 4 or 6 mM) in ACSF, or by the addition of 4-aminopyridine (10 μM) to attain the highest P_r_ (Fig. 1d-f). The M-V analysis revealed significantly higher P_r_ at 2 mM [Ca^2+^]_out_ in warm-cut slices compared to cold-cut slices (P = 0.01, Welch’s *t*-test) (Fig. 1g, Table.S1). The quantal size of synaptic transmission estimated from the MV analysis showed no significant differences between them (Fig.1g, Table.S1). These results indicate that the temperature during slice preparations influence the machinery of neurotransmitter release but has little effect on the kinetics of postsynaptic events.

### Dendritic spine density of Purkinje cells in cerebellar slices prepared at ice-cold and physiological temperature

It has been reported that the exposure of hippocampal slices to CT causes spine loss and beading of the dendrites, and re-warming of the chilled slices induces proliferation of spine structures^8^. Although these structural changes can be minimized by the replacement of NaCl in the cutting solution to sucrose^8^, the risk of the ultrastructural reorganization of spines remains.

Since the loss and proliferation of postsynaptic dendritic spines were prevented by slicing of the hippocampus at room temperature^14^, we hypothesized that slicing of the cerebellum at PT may also prevent the potential reorganization of PC spines. To address this point, we examined differences in density of spines along PC dendrites in acute cerebellar slices prepared at CT and PT. Since the spine density of PC dendrites is too high to measure accurately by conventional confocal microscopy (6-7 spines/μm)^15,16^, we examined it by acquiring z-stacks of tissue volumes with resolution increase in all three spatial directions (3D) by stimulated emission depletion (STED) microscopy. To image isolated single PC dendrites for estimating the spine densities, we used the MADM-11 mouse model expressing green fluorescence protein (GFP) in a small subset of PCs^17^.

The acute slices (parasagittal, 200 μm thickness) were immersion-fixed immediately after slicing (0 h) or after 1-h recovery time in normal ACSF at 37 °C (1 h). PCs were immunolabeled with an anti-GFP antibody and a secondary antibody combined with a dye suitable for high-performance STED imaging (Abberior STAR RED) (Fig.2a-c). The density of dendritic spines immediately after slicing was significantly lower by 40% in the cold-cut slices than that in the perfusion-fixed tissue (P < 0.01) (Fig.2d, Table.S2). After 1-h incubation at 37 °C for recovery, the spine density increased and reached the same level as observed in the perfusion-fixed tissue. In contrast, the spine density in the warm-cut slices showed no significant difference compared to that in the perfusion-fixed tissue and remained unchanged through the recovery time. These observations suggest substantial reorganization of the dendritic spines during slicing at CT and recovery at 37 °C. In contrast, the spine structure can be kept stable in the warm-cutting method.

**Fig. 2.**
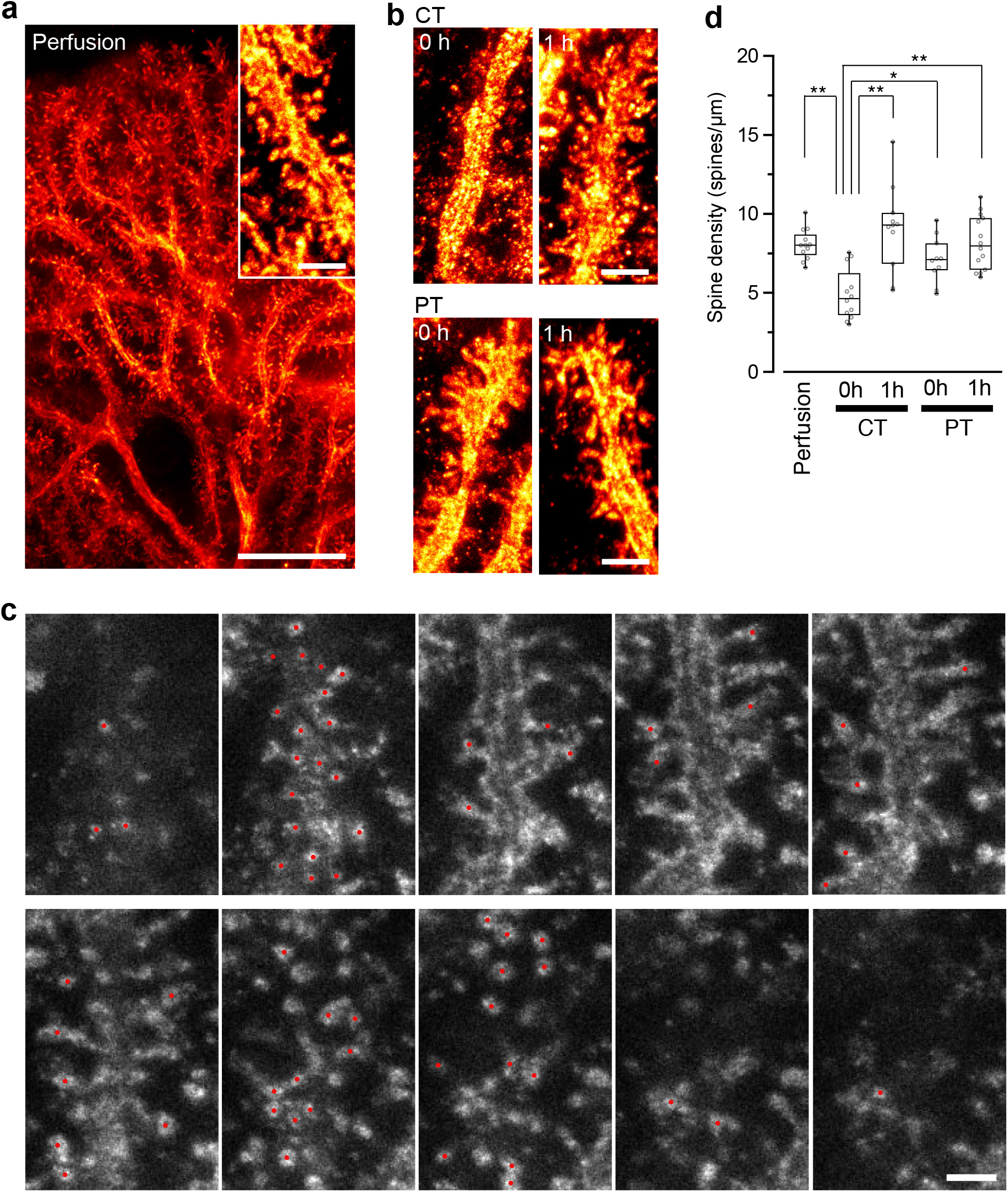
Spine density of Purkinje cells in acute cerebellar slices prepared at ice-cold and physiological temperatures. **a**, Representative super-resolution STED image of PC dendrites and spines (inset) in perfusion-fixed cerebellar tissue. Scale bar = 10 μm and 2 μm (inset). **b**, Representative STED images of spines along a dendrite of PC in acute cerebellar slices immediately after slicing (0 h) or after 1-h recovery (1 h). The images show maximum projections of 20 stacks or 1.8 μm. Scale bar = 2 μm. **c**, Counting dendritic spines observed with 3D STED microscope. Representative z-stack images of a PC dendrite with spines. Red circles indicate individual spines. Scale bar = 1 μm. Optical sections are spaced by 90 nm in the z-direction. **d**, Summary of the dendritic spine density of PCs in perfusion-fixed, cold-cut and warm-cut slice preparations. Each scatter indicates a value obtained from an individual dendrite (perfusion-fixed: n = 12 dendrites, CT/0 h: n = 12 dendrites, CT/1 h: n = 11 dendrites, PT/0 h: n = 10 dendrites, PT/1 h: n = 14 dendrites). Asterisks indicate significant differences (*P < 0.05, **P < 0.01, one-way ANOVA with *post-hoc* Tukey-Kramer test).

### Synaptic vesicle distribution in presynaptic boutons of parallel fibers in cerebellar slices

Synaptic vesicles (SVs) at nerve terminals are accumulated at active zones (AZs) for rapid neurotransmitter release. Cytoskeleton including actin filament and microtubules contributes SV gathering and mobility^18,19^. The dynamics of the cytoskeleton induced by their polymerization/depolymerization is highly temperature-dependent, and exposure of the cytoskeleton to cold temperature disrupts actin filaments and microtubules^20^. These observations indicate a possibility that exposure of brain slices to cold temperature causes rearrangement of SVs in nerve terminals. To address this, we next observed SV distributions in serial ultrathin sections (40 nm thick) by EM in the presynaptic PF boutons in the acute cerebellar slices (parasagittal, 200 μm thick) prepared at CT or PT (Fig.3a). The area of the presynaptic AZs defined as the membrane facing postsynaptic density (PSD) was not significantly different between perfusion-fixed tissues and acute slices prepared at CT and PT, and between slices immersion-fixed immediately after slicing (0 h) and after 1h-incubation at 37 °C (1h) (Supplementary Fig.1a). The numbers of docked SVs (dSVs) near AZs membrane within 5 nm (Fig.3b) were linearly correlated with the AZ areas in all conditions (Supplementary Fig.1b). The total SV numbers were also not significantly different between these preparations (Fig.3c, left). In acute slices at 0 h, the number of dSVs showed no significant difference between perfusion-fixed tissues and slices prepared at CT or PT (Fig.3c, middle), though their density in slices prepared at PT was significantly higher than that in the perfusion-fixed tissues (Table.S3, Fig.3c, right). At 1h, the dSV number and density in the cold-cut slices significantly increased and reached higher levels than those in perfusion-fixed tissues (Table.S3, Fig.3c, middle and right). The density in the cold-cut slices after 1-h recovery was significantly larger compared to the other preparations (Table.S3). In contrast, the docked SV number and density in the warm-cut slices showed no significant changes during the incubation at 37 °C. These results suggest that the SV distribution in nerve terminals is reorganized in the cold-cutting method, while the warm-cutting method can keep the SV distribution stable during acute slice preparation.

**Fig. 3.**
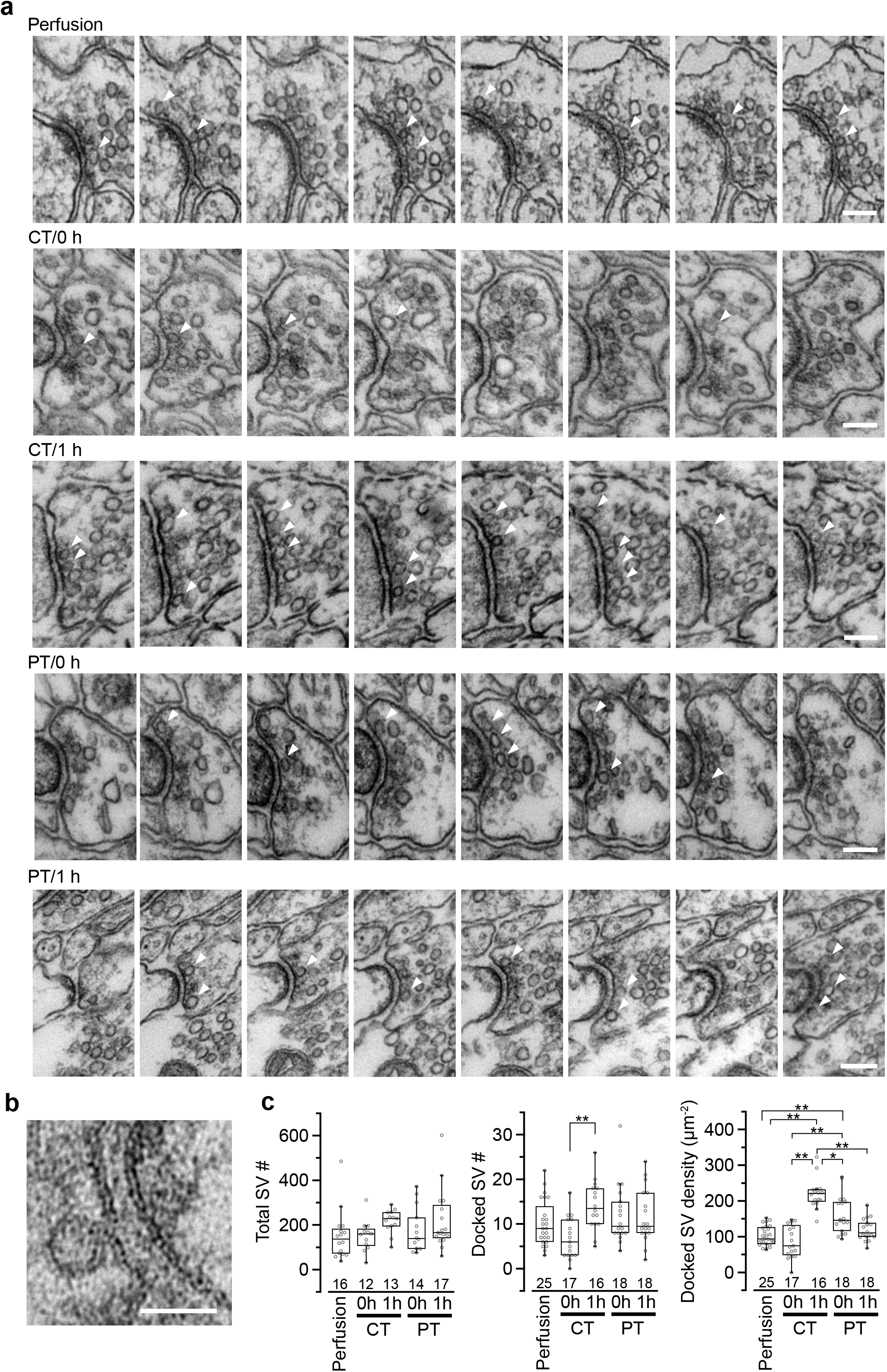
Synaptic vesicle distribution in parallel fiber boutons on PF-PC synapses. **a**, Example images of serial ultrathin sections (40-nm interval) of PF boutons in perfusion-fixed tissue and immersion-fixed acute cerebellar slices with (1 h) or without (0 h) 1-h recovery time. Arrowheads indicate dSVs. Scale bar = 100 nm. **b**, An example image of a dSV, which is located within 5 nm from the AZ membrane in the perfusion-fixed tissue. Scale bar = 50 nm. **c**, Summary of the total SV number in presynaptic boutons (left), the dSV number (middle) and density (right) at AZs. Each scatter indicates a value obtained from an individual bouton. Numerals in plot indicate the numbers of analyzed boutons for each group. Asterisks indicate significant differences (*P < 0.05, **P < 0.01, Kruskal-Wallis H test with *post-hoc* Mann-Whitney *U*-test with Bonferroni correction).

### Pre- and postsynaptic protein distribution in PF-PC synapses

It has been reported that rat hippocampal slices prepared at CT have reduced protein levels for AMPARs, especially GluA1 and GluA3 subunits after incubation at 37 °C^9^. Besides, the cytoskeleton including actin filaments works as an anchor and/or a trafficking pathway for membrane-associated proteins and regulates the distribution of proteins including AMPARs at the postsynaptic membrane^21^. These reports indicate a possibility that the amount and distribution of membrane-associated proteins at pre- and postsynaptic sites are altered through the brain slice preparation at CT. To investigate the two-dimensional distribution of synaptic proteins contributing synaptic transmission in the AZ and postsynaptic area of PF-PC synapses, we performed SDS-digested freeze-fracture replica labeling (SDS-FRL) for AMPAR (GluA1-3), GluD2, RIM1/2, and Cav2.1 (Fig.4). The cerebellar slices prepared at CT and PT (parasagittal, 200 μm thick) were immersion-fixed immediately after slicing (0 h) or after the 1-h recovery in ACSF at 37 °C (1 h) and were frozen using high pressure freezing machine for fracturing. Postsynaptic areas on the exoplasmic face (E-face) and presynaptic AZs on the protoplasmic face (P-face) in freeze-fracture replica samples were identified with aggregation of intramembrane particles at the electron microscopic level as described previously^22–24^. To identify PF-PC synapses in replica samples of the molecular layer, labeling for GluD2 or vesicular glutamate transporter 1 (VGluT1) were used as markers^24,25^. Immunogold particles for GluA1-3 were highly concentrated in the postsynaptic areas labeled for GluD2, while those for RIM1/2 and Cav2.1 were concentrated in the AZs of VGluT1-labeled presynaptic profiles as previously described^24,26^ (Fig.4a).

**Fig. 4.**
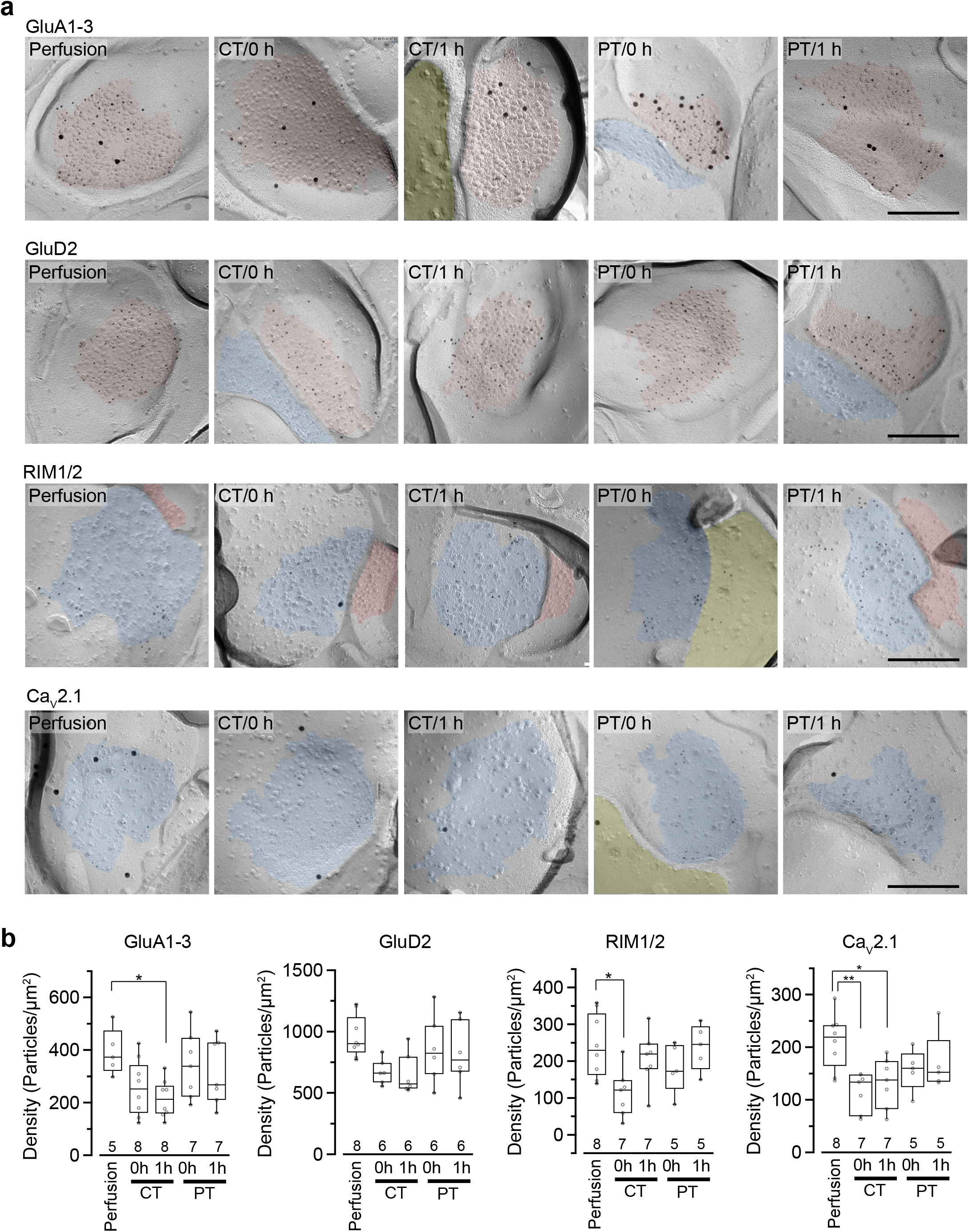
Distribution of synaptic proteins in PSD and AZs. **a**, Representative images of freeze-fracture replicas of PF-PC synapses labeled for GluA1-3, GluD2, RIM1/2 and Cav2.1 (5-nm gold) with PF-PC markers (GluD2 for GluA1-3, VGluT1 for RIM1/2 and Cav2.1, 15-nm gold). PSD on the exoplasmic face (E-face), AZs on the protoplasmic face (P-face) and cross-fractured cytoplasm were indicated with red, blue and yellow, respectively. Scale bar = 200 nm. **b**, Summary of gold particle density for GluA1-3, GluD2, RIM1/2, and Cav2.1 on PSD or AZs. Each scatter indicates the mean value obtained from an individual replica. Numerals in plot indicate the numbers of analyzed replicas for each group. Asterisks indicate significant differences (*P < 0.05, **P < 0.01, one-way ANOVA with *post-hoc* Tukey-Kramer test).

For the quantitative analysis of protein distributions on pre- and postsynaptic membranes, we measured the density of immunogold particles for each protein within the synaptic areas (Fig.4b, Table.S4). AT 0h, labeling for GluA1-3 showed no significant difference between preparations including perfusion-fixed tissues. However, after the recovery time, the GluA1-3 labeling in slices prepared at CT was significantly lower than that in the perfusion-fixed tissue (P < 0.05) (Table.S4, Fig.4b). In contrast, GluA1-3 labeling in the slices prepared at PT was similar to that in perfusion-fixed tissues both at 0 h and 1 h. The labeling for GluD2 showed no significant differences between these preparations. For RIM1/2 at AZs, the labeling density in the slices prepared at CT temporarily decreased after slicing and recovered during the recovery time. The RIM1/2 labeling in the warm-cut slices was similar to that in perfusion-fixed tissue and kept constant during the recovery. For Cav2.1 at AZs, the labeling in cold-cut slices was significantly lower than that in the perfusion-fixed tissue (Table.S4, Fig.4b) and only partially recovered after the 1-h recovery time, whereas the warm-cut slices showed no significant differences in Cav2.1 labeling between these preparations (Fig.4b). These results indicate that the exposure of the brains to cold temperature alters synaptic protein distribution at both pre- and postsynaptic sites, and some of these changes do not recover by the incubation of slices at 37 °C for 1 h. The warm-cutting method can circumvent these problems.

### Long-term plasticity induced by HOKR adaptation

The changes of synaptic properties during recovery time may affect the measurement of synaptic plasticity occured *in vivo*. Thus, we finally tested cold- and warm-cutting methods for detectability of LTD induced *in vivo* using a simple model of cerebellar motor learning. LTD of PF–PC synapses is associated with decreases in the number of synaptic AMPARs^16^ and reported to be a primary cellular mechanism of adaptation of horizontal optokinetic response (HOKR) in mouse^27^. One hour HOKR training induced a transient AMPAR reduction by 28% in PF-PC synapses in the flocculus^16^, which controls horizontal eye movement^28^. This result predicts a reduced EPSCs in PF-PC synapses after HOKR adaptation. Indeed, a further study using conventional acute slice preparation showed the decrease in quantal EPSC amplitude of PF-PC synapses in the flocculus of mice after HOKR training, but the reduction was below 10%, much lower than the value expected from the reduction of AMPAR density^29^.

Our study of SDS-FRL showed that the AMPAR density on postsynaptic sites in acute cerebellar slice prepared at CT but not PT was significantly lower after the recovery than that in the perfusion-fixed preparation (Fig.4b). This result could indicate a higher sensitivity in warm-cut slices to detect the quantal EPSC amplitude changes in the flocculus induced by the HOKR training. To examine this possibility, we recorded spontaneous miniature EPSC (mEPSC) from PCs in the flocculus after 1-h HOKR training. Continuous horizontal optokinetic stimulation enhanced the eye movement of the trained mice, and the gain of HOKR, which was defined as the amplitude of eye movement divided by the screen movement significantly increased by 62% (1 min: 0.54 [0.43-0.56], n = 10; 60 min: 0.78 [0.75-0.80], n = 10, P < 0.01) (Fig.5a). We sacrificed the trained mice within 1 min after the training, and sliced the cerebellum (coronal, 250-300 μm thick) in the cutting solution at CT or PT within 30 min after the decapitation. Then we used the slices for mEPSC recordings immediately after slicing (warm-cut slices) or after the 1-h recovery at 37 °C in normal ACSF (cold-cut slices) (Fig.5b). The mean amplitude of mEPSC in control was not significantly different between the cold-cut and the warm-cut slices, but the amplitude in cold-cut slices varied more than that in warm-cut slices (CV: 0.24 for CT, 0.11 for PT, Fig.5e-f). The frequency of spontaneous mEPSCs recorded from PCs in the flocculus was not significantly changed by the HOKR training in both cold- and warm-cut slices (Fig.5b). The mEPSC frequency in the cold-cut slices was significantly higher than that in the warm-cut slices for both control and trained animals (Fig.5e-f, Table.S5). After the 1-h HOKR training, mEPSC amplitude was significantly decreased by 23.8% in the warm-cut slices (Fig.5d and f, Table.S5), consistent with our previous finding of decrease in AMPARs by 28%^16^. The amplitude in the cold-cut slices also showed a tendency to decrease but the difference did not reach statistical significance (Fig. 5c and e, Table.S5). The higher detectability of LTD in the warm-cut slices compared to cold-cut slices indicates an advantage of the warm-cutting method for investigating synaptic plasticity induced by behavior experiments.

**Fig. 5.**
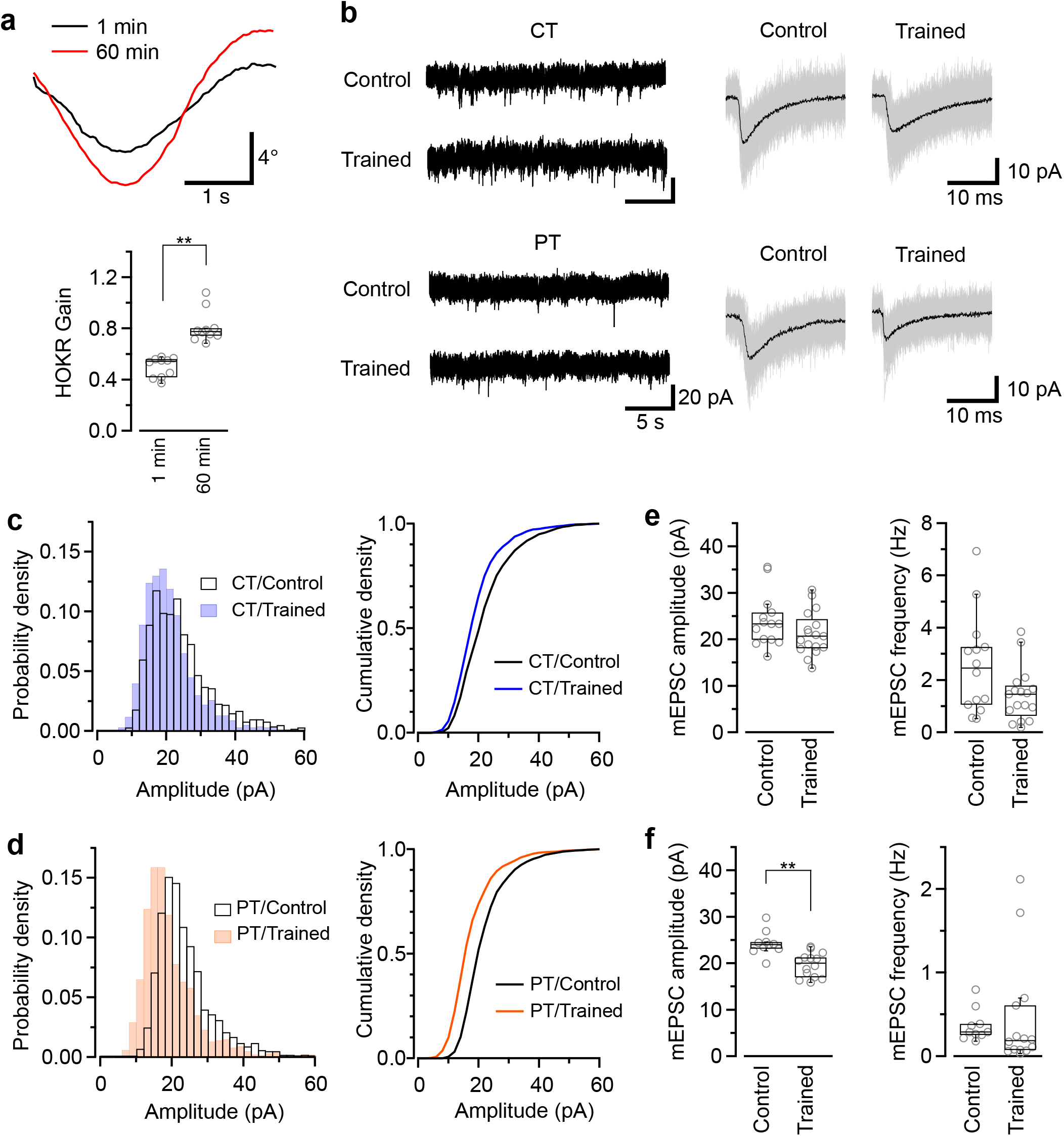
Application of warm-cutting slice preparation method to the detection of long-term depression by HOKR training. **a**, HOKR adaptation. Top, representative eye-movement traces of a mouse before and after 1-h HOKR training. Bottom, HOKR gain changes induced by 1-h HOKR training (P < 0.01, paired *t*-test). Each scatter indicates the mean value obtained from individual animal (n = 10 animals for each). **b**, Representative traces of spontaneous mEPSC events recorded from PCs at cerebellar flocculus of untrained (control) and trained mice. Right, superimposed mEPSC traces (50 events, gray). and average of the events (black traces). **c-d**, Histogram (left) and cumulative curve (right) of mEPSC amplitude distribution recorded from PCs in cold-cut (**c**) and warm-cut (**d**) slices of control and HOKR-trained mice. **e-f**, Box plot of mEPSC amplitudes (left) and frequency (right) recorded from PCs in cold-cut (**e**) and warm-cut (**f**) slices of control and HOKR-trained mice. Each scatter indicates the mean amplitude and frequency of mEPSCs obtained from an individual PC (CT/control: n = 14 cells, CT/trained: n = 17 cells, PT/control: n = 10 cells, PT/trained: n = 14 cells). Asterisks indicate significant differences (**P < 0.01, Welch’s *t*-test).

## Discussion

In this study, we show that brain slice preparation at PT better preserves molecular and ultrastructural properties of synapses compared to the conventional slice preparation at CT. These results suggest strong advantages of warm-cutting method for investigating synaptic functions.

### Acute brain slice preparation at PT

Here we prepared mouse cerebellar slices at CT and PT and compared the synaptic properties in these tissues. The brain in cutting solution at PT was softer than that at CT, so we needed to adjust the parameters of the slicers for the warm-cutting method (see Methods). The parameters may need to be adjusted for each brain region, age and also species of animals.

### Preservation of synaptic properties in acute brain slices prepared at PT

To investigate synaptic functions in brains, synaptic properties in acute brain slices should be kept as close as those in the intact brain. In the present study, AMPAR and Cav2.1 in the cold but not warm-cut slices showed significantly lower densities than those in the perfusion-fixed tissue. Although some of the structural parameters of the synapse in perfusion-fixed tissue can be altered by formaldehyde fixation^30,31^, we used the same fixative, temperature and time of fixation to compare these properties between perfusion-fixed tissues and immersion-fixed acute slices. Assuming similar effects of chemical fixation between these preparations, our study suggest that the warm-cutting method preserves molecular and structural properties of synapses better than the conventional cold-cutting method. We also found that PF-PC synapses in the cold-cut slices have higher mEPSC frequency than that in warm-cut slices. Although larger P_r_ may increase the frequency of mEPSCs, P_r_ measured by M-V analysis in cold-cut slices is significantly smaller than that in warm-cut slices, which is consistent with the reduced Cav2.1 density at AZs in cold-cut slices. Readily releasable pool (RRP) size of SVs also correlates with mEPSC frequency, and the number of dSVs may indicate RRP size^32,33^. Thus, the increased number and density of dSVs after the recovery in cold-cut slices could contribute to the higher mEPSC frequency. Another possibility is instability of molecular machinery for neurotransmitter release in cold-cut slices contributing to the higher frequency of mESPCs. Although functional properties of synapses in truly intact brains *in vivo* are mostly unknown, our findings indicate that brain slices prepared at PT have an advantage in preserving the synaptic function compared to those prepared at CT.

### Changes of molecular and structural properties of synapses during recovery time in cold-cut brain slices

Another advantage of the warm-cutting method is the stability of synaptic properties during the slice preparation. The PC spine density values obtained with 3D STED microscopy are comparable with those obtained by high-voltage EM^16^ and 3D EM reconstruction analysis^15^, indicating the reliability of our data. We found that 40% of spines along PC dendrites disappeared after slicing at CT and then recovered after 1-h recovery time, consistent with previous studies on the spine reorganization in acute hippocampal slices^7,8^. Brain slicing at CT also modifies SV distributions in synaptic boutons and the density of RIM1/2, a SV-associated protein at AZs, which recover after 1-h incubation time at 37 °C. In contrast, cerebellar slices prepared at PT can keep these post- and presynaptic features stable throughout the slice preparation. Consequently, brain slices prepared at PT can be used immediately after slicing for the electrophysiological experiments as a “ready-to-use” slice preparation. This is a major advantage especially for investigating synaptic plasticity induced in behavioral experiments as discussed below.

### Application of warm-cutting method for investigating synaptic plasticity induced by behavior experiments

In acute slices prepared at PT, the reduction of mEPSC amplitude in PCs induced by HOKR training was larger than that in the cold-cut slices. Why is the difference more detectable in warm-cut slices than in cold-cut slices? One possibility is the difference in variation of mEPSC amplitude. The CV value of mEPSC amplitude in cold-cut slices was larger than in warm-cut slices, which could be due to high variability of quantal size in the reconstructed spine synapses. Another possibility is that LTD in synapses induced by the training might be partially reset through the spine reconstruction after slicing at CT. AMPAR density is regulated by various proteins including enzymes (e.g. CaMKII and PP2A) and actin^34,35^, and exposure to cold temperature reduces metabolic activity of these enzymes and also depolymerizes actin filaments^20^. Cooling and rewarming of brain tissues might reset the activities of these proteins reorganizing AMPAR distribution on postsynaptic membrane and obscure the plastic changes occurred *in vivo*. Lastly, earlier timing of measurements with the warm-cut slices after the training *in vivo* could contribute to the better detection of synaptic plasticity lasting less than a few hours.

In summary, we demonstrate that the warm-cutting method has substantial advantages over the conventional cold-cutting method, e.g. highly preserved synaptic properties, stability of synaptic properties through the preparing processes, and “ready-to-use” acute slice preparation, for investigating synaptic functions. These advantages should facilitate a wide range of neuroscience research, especially for synaptic plasticity induced *in vivo*.

## Materials and Methods

### Animals

Animal experiments were conducted in accordance with the guideline of Institute of Science and Technology Austria (IST Austria) (Animal license number: BMWFW-66.018/0012-WF/V/3b/2016). Mice were bred and maintained in the Preclinical Facility of IST Austria. Unless otherwise noted, C57BL/6J mice of either sex at postnatal (P) 4-6 weeks were used in this study.

### Acute brain slice preparation

Mice were decapitated under isoflurane anesthesia and their brains were quickly removed from the skull and immersed into cutting solution contained (in mM): 300 sucrose, 2.5 KCl, 10 glucose, 1.25 NaH_2_PO_4_, 2 Na Pyruvate, 3 *myo*-inositol, 0.5 Na ascorbate, 26 NaHCO_3_, 0.1 CaCl_2_, 6 MgCl_2_ (pH 7.4 when gassed with 95% O_2_/5% CO_2_) at ice-cold temperature (CT, < 4 °C) or physiological temperature (PT, 35-37 °C). The cerebellum was dissected from the whole brain and immediately glued on a cutting stage of a tissue slicer (Linear Slicer Pro7, Dosaka EM, Kyoto, Japan). The parameters of the slicer were optimized for slicing at each temperature (CT: amplitude = 4.5, frequency = 84-86 Hz, speed = 2.0-2.5, PT: amplitude = 5.5, frequency = 84-86 Hz, speed = 2.0). Bath temperature was kept within the desired range (below 4 °C for CT, 35-37 °C for PT) by adding cold water with crushed ice or warm water into the bath of the slicer and was monitored throughout the cutting procedure with a thermometer. Slices were then maintained in the standard ACSF contained (in mM): 125 NaCl, 2.5 KCl, 10 glucose, 1.25 NaH_2_PO_4_, 2 sodium pyruvate, 3 *myo*inositol, 0.5 sodium ascorbate, 26 NaHCO_3_, 2 CaCl_2_, 1 MgCl_2_ (pH 7.4 when gassed with 95% O_2_/5% CO_2_) at 37 °C for 1 hour and subsequently at room temperature (RT) until use.

### Fixations

We used three types of fixative solutions; 2.5% glutaraldehyde and 2% paraformaldehyde (PFA) in 0.1 M HEPES buffer (pH 7.4 adjusted with NaOH) for serial ultrathin sectioning with electron microscopic analysis, 2% PFA and 15% picric acid in 0.1M sodium phosphate buffer (s-PB, pH 7.4) for SDS-FRL, and 4% PFA and 0.05% glutaraldehyde in 0.1 M s-PB for immunofluorescence imaging with a STED microscope.

For the perfusion-fixation of brains, the mice were anesthetized with ketamine/xylazine mixture via intraperitoneal injection and perfused transcardially with 0.1 M HEPES buffer (for serial sectioning) or 0.1 M s-PB (for SDS-FRL and STED imaging) for 1 min, followed by the fixative solution for 12 min. The perfusion of the buffers and fixatives was done at room temperature (23-25 °C). The brains were then removed from the skull and post-fixed for 1-2 h at RT and then kept in the fixative solution overnight at 4 °C. Sagittal slices of the cerebellum were cut using a tissue slicer (Linear Slicer Pro7, Dosaka, Kyoto, Japan) in 0.1 M HEPES buffer or 0.1 M s-PB. The acute cerebellar slices were immersion-fixed in the fixative solution for each purpose at RT for 1-2 h on a shaker (160 rpm) and then kept in the fixative solution overnight at 4 °C.

### Super-resolution fluorescence imaging with STED microscope

For super-resolution imaging with the STED microscope, we used Mosaic Analysis of Double Marker (MADM-11) mice crossed with ubiquitous Cre driver (Hprt-cre) mice^17^. The fixed cerebellar slices (parasagittal, 200 μm thick) of MADM-11 mice (P5-7w, males) were washed three times with PBS and blocked in blocking buffer (3% BSA + 0.1% Triton X-100 in PBS) for 1-2 hr at RT. The slices were then incubated with the monoclonal mouse IgG antibody against GFP (1 μg/ml, Abcam) dissolved in the blocking buffer overnight at 4 °C. The slices were washed three times with PBS and incubated with the secondary antibody against mouse IgG conjugated with STAR RED fluorescent dye (1:100, goat IgG, Abberior) dissolved in the blocking buffer overnight at 4 °C. The slices were washed three times with PBS, transferred on the slide, covered with mounting medium (Abberior Mount Liquid Antifade) using coverslip (#1.5) and sealed with nail polish.

3D STED microscopy was performed on a commercial inverted STED microscope (Expert Line, Abberior Instruments, Germany) with pulsed excitation and STED lasers. A 640 nm laser was used for excitation and a 775 nm laser for stimulated emission. A silicone oil immersion objective with numerical aperture 1.35 (UPLSAPO 100XS, Olympus, Japan) and a correction collar was used for image acquisition. The fluorescence signal was collected in a confocal arrangement with a pinhole size of 0.8 Airy units using a photon counting avalanche photodiode with a 685/80 nm bandpass filter for STAR RED detection. The pulse repetition rate was 40 MHz and fluorescence detection was time-gated. The imaging parameters used were 20 μs pixel dwell time and two line accumulations. Laser powers at the sample were 2-3 μW (640 nm) excitation and 20-30 mW STED laser power, the power ratio in the xy-STED “doughnut” beam and the beam providing additional resolution increase in the axial (*z*)-direction (“z-doughnut”) was between 40/60 and 60/40. Voxel size was 30 x 30 x 90 nm. Stacks typically spanned 5 μm in z-direction, covering the full dendrite. We observed the stained dendrites within tens of micrometer from the surface of the tissues. Individual dendritic spines of PCs were manually counted using FIJI software (distributed under the General Public License).

### Serial ultrathin sectioning with electron microscopy

The fixed cerebellar slices (parasagittal, 200 μm thick) were washed three times with 0.1M HEPES buffer and then treated with 1% osmium tetroxide (30 min at RT) and 1% uranyl acetate (30 min at RT) to provide adequate contrast for electron microscopic analysis. Following the dehydration with a series concentration of ethanol (50, 70, 90, 96 and 100%) and acetone, the slices were infiltrated with and embedded in Durcupan resin at 60 °C for 3 days. The cerebellar cortex including the molecular layer at lobule V-VI was trimmed and exposed, and serial sections of 40-nm thickness were obtained (within a few microns from the surface of tissues) using an ultramicrotome (UC7, Leica). The serial sections were picked on the grid coated with pioloform and counterstained with 1% uranyl acetate (7 min at RT) and 0.3% lead citrate (4 min at RT). Serial images were taken from the samples under a transmission electron microscope (TEM, Tecnai 10, FEI) with iTEM (OSIS) or RADIUS (EMSIS). The images were analyzed using Reconstruct (SynapseWeb, Kristen M. Harris, PI) and FIJI software. Thickness of the ultrathin sections measured by minimal folds method^36^ was 40.3 nm (35.6-46.8 nm, n = 8). Active zones (AZs) are defined as the membrane facing postsynaptic density (PSD) which is clearly identified as an electron-dense thickening in dendritic spines. Docked synaptic vesicles (dSVs) were defined by the distance from AZ membranes (< 5 nm, Fig.3b).

### Antibodies

Rabbit polyclonal antibodies against AMPA receptor subunits were raised against synthetic peptides with the following sequences: anti-GluA1-3, (C)VNLAVLKLSEQGVLDKLKSKWWYDKGE (residues 760-786 of mouse GluA1), anti-GluA1, (C)SMSHSSGMPLGATGL (residues 893-907 in the C-terminal intracellular region of rat GluA1),anti-GluA3, (C)NEYERFVPFSDQQIS (residues 394-408 in the N-terminal extracellular region of rat GluA3), and anti-GluA4, (C)GTAIRQSSGLAVIASDLP (residues 885-902 in the C-terminal intracellular region of rat GluA4). Mouse monoclonal anti-GluA2 was raised against the synthetic peptide (C)YKEGYNVYGIESVKI (residues 869-883 in the C-terminal intracellular region of rat GluA2). Affinity-purified antibodies were obtained from antisera using respective antigen peptides. Mouse anti-actin antibody was obtained from BD Biosciences.

### Plasmids

Full-length rat GluA1 and GluA2 cDNAs were subcloned into pcDNA3 (plasmid vector, Invitrogen) to generate pc3-GluA1 and pc3-GluA2. In addition, full-length murine GluA3 and GluA4 cDNAs were subcloned into pRK5 (plasmid vector) to generate pRK5-GluA3 and pRK5-GluA4.

### Transfection and immunoblot analysis

COS-7 cells were maintained in Dulbecco’s modified Eagle’s medium (DMEM) with 10% fetal bovine serum (FBS) and were transiently transfected with GluA1-4 expression plasmids using Lipofectamine 2000 (Invitrogen). After 48 h of transfection, the cultured cells were harvested and lysed in 1× SDS sample buffer. Cell lysates were separated by SDS-PAGE, followed by electrotransfer into PDVF membrane and incubated with respective antibodies. Protein bands were developed by the enhanced chemiluminescence method using a commercially available kit (Nacalai Chemicals).

### SDS-digested freeze-fracture replica labeling (SDS-FRL)

The fixed cerebellar slices (parasagittal, 150 μm thick) were washed three times with 0.1M PB and then immersed in graded glycerol of 10-20% in 0.1 M PB at RT for 10 min and then 30% glycerol in 0.1M PB at 4 °C overnight for the cryoprotection. The molecular layer of the cerebellum at lobule IV-VI was trimmed from the slices and frozen by a high pressure freezing machine (HPM010, BAL-TEC). The frozen samples were fractured into two parts at −130 °C and replicated by carbon deposition (5 nm thick), carbon-platinum (uni-direction from 60°, 2 nm) and carbon (20-25 nm) in a freeze-fracture machine (JFD-V, JOEL, Tokyo, Japan). The samples were digested with 2.5% SDS solution containing 20 mM sucrose and 15 mM Tris-HCl (pH 8.3) at 80 °C for 17 h. The replicas were washed in the SDS solution, 2.5% bovine serum albumin (BSA) contained SDS solution and then 50-mM Tris-buffered saline (TBS, pH 7.4) containing 5%, 2.5% and 0.1% BSA at RT for 10 min for each. To avoid non-specific binding of antibodies, the replicas were blocked with 5% BSA in TBS for 1-2 h at RT. The replicas were then incubated with the primary antibodies dissolved in TBS with 2% BSA at 15 °C overnight. For the labeling of AMPARs, the rabbit polyclonal antibody for GluA1-3 (4.3 μg/ml) was used with guinea pig polyclonal antibody for GluD2 (0.61 μg/ml, provided by Masahiko Watanabe) used as a marker of PF-PC synapses^16,24,37^. The specificity of the GluA1-3 antibody was checked by Western-blotting (Supplementary Fig.2). For GluD2 labeling, polyclonal guinea pig antibody for GluD2 (4.1 μg/ml) was used ^16,24^ For the labeling of RIM1/2 and Cav2.1 subunit of voltage-gated calcium channel, rabbit polyclonal antibody for RIM1/2 (5.0 μg/ml, Synaptic Systems) and guinea pig polyclonal antibody for Ca_V_2.1 (4.1 μg/ml, provided by Masahiko Watanabe)^38,39^ were used respectively, with mouse monoclonal antibody for VGluT1 (5.0 μg/ml, NeuroMab) as a marker of PF boutons^25^. The replicas were then washed three times with TBS with 0.1% BSA and incubated with the secondary antibodies conjugated with gold particles (5 or 15 nm diameter, goat IgG, BBI) dissolved in TBS with 2% BSA (1:30) overnight at 15 °C. After immunogold labeling, the replicas were washed with TBS with 0.1% BSA (twice) and distilled water (twice) and picked up onto a grid coated with pioloform. Images (20-25 images per replica) were obtained under TEM (Tecnai 10) with iTEM (OSIS) or RADIUS and analyzed with FIJI software to calculate the density of gold particles defined by the particle number divided by PSD/AZ area. PSD and AZs were indicated with the aggregation of intramembrane particles on the replica at the electron microscopic level as described previously^22,23,24^.

### Whole-cell patch-clamp recording

Acute cerebellar slices (lobule IV-V, coronal, 250-300 μm thick) were superfused with oxygenated ACSF in the recording chamber and visualized using upright microscope (BX51WI, Olympus, Japan) with a 20x water-immersion objective lens. Data were acquired at a sampling rate of 50 kHz using an EPC-10 USB double patch-clamp amplifier controlled by PatchMaster software (HEKA, Germany) after online filtering at 5 kHz. Experiments were performed at RT (26-27 °C) within 5 h after slicing at both temperatures. Resistances of patch electrodes were 3–6 MΩ. The series resistances were 6–15 MΩ and compensated for a final value of 5 MΩ.

Throughout the experiments, recordings from Purkinje cells were made in voltage-clamp mode at a holding potential of −70 mV. The recordings were performed up to 1 h after rapturing. The pipette solution contained the following (in mM): 110 Cesium methanesulfonate, 30 CsCl, 10 HEPES, 5 EGTA, 1 MgCl_2_, and 5 QX314-Cl (pH 7.3, adjusted with CsOH). EPSCs were evoked by electrical stimulation (0.5–5.0 V, 60 μs) through an ACSF-filled glass pipette electrode placed on the molecular layer of the cerebellar slices using a stimulus isolator (ISO-Flex, A.M.P.I, Israel). The grass pipette for the stimulation was the same size of the one for recording. The distance between the stimulation and recording electrodes was 200-500 μm to avoid the contamination of climbing fiber inputs or the excitation of recording PCs. The EPSC recordings were performed in the presence of bicuculline methiodide (10 μM) to block GABAergic inhibitory synaptic current. Data was off-line analyzed using Igor Pro 6 software (WaveMatrics, Oregon, USA) with Patcher’s Power Tools (Max-Planck-Institut für biophysikalische Chemie, Göttingen, Germany).

Mean-variance (M-V) analysis was used to estimate the mean release probability per site (P_r_). EPSCs, evoked with a 15-s interval, were recorded in the presence of the low-affinity glutamate receptor antagonist kynurenic acid (2 mM) to reduce possible effects of AMPAR saturation^40^. Under various extracellular Ca^2+^ concentrations ([Ca^2+^]_out_, 1.5-6 mM) for altering Pr, 15 successive EPSCs were collected for constructing an M-V plot. To acquire data at highest P_r_, the K^+^ channel blocker 4-aminopyridine (10 μM) was added to the ACSF. M-V plots were analyzed by fitting the simple parabola equation:

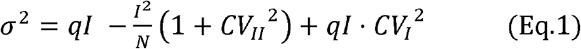

where I and σ^2^ represent the mean amplitude and variance of EPSCs, respectively, q denotes mean quantal size and N denotes the number of release sites. CV_I_ and CV_II_ mean the coefficients of intrasite and intersite quantal variability respectively, assumed to be 0.3^12^. In this method, assuming that σ^2^ arises entirely from stochastic changes in P_r_, q can be estimated from the initial slope of the parabola, Nq from the larger X intercept of the parabola, and P_r_ can be estimated as I/Nq.

### Horizontal optokinetic response (HOKR) training and miniature EPSC recording

HOKR was recorded as described previously^25^. C56BL/6J mice (P8-11w, male) were implanted with a 15-mm-long bolt on the skull with a synthetic resin under isoflurane anesthesia and allowed to recover at least for 24 h. The mouse was mounted on the turntable surrounded by a checked-pattern screen with the head fixed via the bolt, and its body was loosely restrained in a plastic cylinder. The frontal view of the right eye, under the illumination of infrared (wavelength, 860 nm) LED, was captured using a vertically positioned CCD camera (SSC-M350, Sony, Japan) and displayed on a 12-inch TV monitor (magnification, 55×). The area of the pupil was determined from the difference in brightness between the pupil and the iris. The real-time position of the eye was measured by calculating the central position of the left and right margins of the pupil at 50 Hz using a position-analyzing system (C-1170; Hamamatsu Photonics, Japan) and stored on a personal computer. HOKR was evoked with sinusoidal screen oscillation at 17° and 0.25 Hz (maximum screen velocity, 7.9° per second) in the light, and its gain was defined as the averaged amplitudes of eye movements vs. those of screen oscillation. The mice were trained for 1 h with sustained screen oscillation. After the training, mice were sacrificed under isoflurane anesthesia and removed their brains from the skull. The cerebellum was dissected from the brain and mounted in 3% low-melting-point agarose (for ice-cold) or 1% agarose (for PT) in Tyrode’s solution contained (mM): 138 NaCl, 3 KCl, 10 HEPES and 7 MgCl_2_ (pH = 7.4, adjusted with NaOH). Acute cerebellar slices containing flocculus/paraflocculus were prepared (coronal, 250-300 μm thick) in cutting solution at CT or PT. The cold-cut slices were pre-incubated in standard ACSF at 37 °C before recording. The warm-cut slices were immediately used for patch-clamp recording without recovery step in ACSF.

Spontaneous miniature EPSC (mEPSC) events were recorded with the same procedure for the evoked EPSC recording as described above. The ACSF during the recordings contained 4 mM CaCl_2_, 0.5 mM MgCl_2_ to increase mEPSC frequency, and also 1 μM tetrodotoxin and 10 μM bicuculline methiodide to block firing and inhibitory synaptic currents. Data were analyzed off-line using Python-based software Stimfit^41^. Spontaneous mEPSCs were detected using a sliding template method^42^ implemented in Stimfit. The templates were made by the averaging of the initial 50-100 mEPSCs selected by eye. Overlapped events were excluded from analysis by visual inspection. The values of the mean mEPSC amplitude for each cell were calculated from the average of 50-200 (PT) or 100-400 (CT) mEPSC events.

### Statistical Analysis

Unless stated otherwise, all data were presented as median (interquartile range [IQR]). The box plots were drawn with whiskers at farthest points within 1.5 x IQR. Normality of data was tested with Kolmogorov-Smirnov test. Normally distributed data were compared with two-tailed Welch’s *t*-test or paired *t*-test for 2 groups. For more than 2 groups, normally distributed data were compared with ANOVA with *post-hoc* Tukey-Kramer test, and others were compared with Kruskal-Wallis H test with *post-hoc* Mann-Whitney *U* test with Bonferroni correction. Comparison is detailed in the respective Results section and Figure Legends. Statistical significance was defined as *p < 0.05 and **p < 0.01. Statistics were performed with Excel (Welch’s *t*-test) and R software (other tests). All P values are shown in supplemental tables.

## Acknowledgments

We thank Peter Jonas for critical comments on the manuscript, Simon Hippenmeyer for providing MADM-11 mice and Masahiko Watanabe for providing GluD2 and Cav2.1 antibodies. This work was financially supported by funding from the European Union’s Horizon 2020 research and innovation program under the Marie Sklodowska-Curie grant agreement No. 793482 (to K.E.) and the European Research Council (ERC) grant agreement No. 694539 (to RS), and Austrian Science Fund (FWF; I 3600-B27, to J.G.D. and P.V.).

## Author contributions

K.E. and R.S. initiated the project. K.E. conducted electrophysiological and electron microscopic experiments. P.V. performed STED microscopic experiments. E.H. performed HOKR training for mice. M.I. and Y.F. generated antibodies against GluA1-3. All authors analyzed data. K.E. and R.S. wrote the paper with comments from all authors.

## Data availability

The data that support the findings of this study are available within the article and its Supplementary Information or from the corresponding author upon reasonable request.

## References

1. Li, C. L. & McIlwain, H. Maintenance of resting membrane potentials in slices of mammalian cerebral cortex and other tissues in vitro. J. Physiol. 139, 178–190 (1957).

2. Yamamoto, C. & McIlwain, H. Electrical activities in thin sections from the mammalian brain maintained in chemically-defined media in vitro. J. Neurochem. 13, 1333–1343 (1966).

3. Yamamoto, C. Recording of electrical activity from microscopically identified neurons of the mammalian brain. Experientia 31, 309–311 (1975).

4. Takahashi, T. Intracellular recording from visually identified motoneurons in rat spinal cord slices. Proc. R. Soc. Lond. 202, 417–421 (1978).

5. Llinás, R. & Sugimori, M. Electrophysiological properties of in vitro Purkinje cell dendrites in mammalian cerebellar slices. J. Physiol. 305, 197–213 (1980).

6. Bischofberger, J., Engel, D., Li, L., Geiger, J. R. P. & Jonas, P. Patch-clamp recording from mossy fiber terminals in hippocampal slices. Nat. Protoc. 1, 2075–2081 (2006).

7. Fiala, J. C. et al. Timing of neuronal and glial ultrastructure disruption during brain slice preparation and recovery in vitro. J. Comp. Neurol. 465, 90–103 (2003).

8. Kirov, S. A., Petrak, L. J., Fiala, J. C. & Harris, K. M. Dendritic spines disappear with chilling but proliferate excessively upon rewarming of mature hippocampus. Neuroscience 127, 69–80 (2004).

9. Taubenfeld, S. M., Stevens, K. A., Pollonini, G., Ruggiero, J. & Alberini, C. M. Profound molecular changes following hippocampal slice preparation: Loss of AMPA receptor subunits and uncoupled mRNA/protein expression. J. Neurochem. 81, 1348–1360 (2002).

10. Huang, S. & Uusisaari, M. Y. Physiological temperature during brain slicing enhances the quality of acute slice preparations. Front. Cell. Neurosci. 7, 1–8 (2013).

11. Fried, N. T., Moffat, C., Seifert, E. L. & Oshinsky, M. L. Functional mitochondrial analysis in acute brain sections from adult rats reveals mitochondrial dysfunction in a rat model of migraine. AJP: Cell Physiology 307, C1017–C1030 (2014).

12. Clements, J. D. & Silver, R. A. Unveiling synaptic plasticity: A new graphical and analytical approach. Trends Neurosci. 23, 105–113 (2000).

13. Scheuss, V., Schneggenburger, R. & Neher, E. Separation of presynaptic and postsynaptic contributions to depression by covariance analysis of successive EPSCs at the calyx of Held synapse. J. Neurosci. 22, 728–739 (2002).

14. Bourne, J. N., Kirov, S. A., Sorra, K. E. & Harris, K. M. Warmer preparation of hippocampal slices prevents synapse proliferation that might obscure LTP-related structural plasticity. Neuropharmacology 52, 55–59 (2007).

15. Ichikawa, R. et al. Distal Extension of Climbing Fiber Territory and Multiple Innervation Caused by Aberrant Wiring to Adjacent Spiny Branchlets in Cerebellar Purkinje Cells Lacking Glutamate Receptor δ2. The Journal of Neuroscience 22, 8487–8503 (2002).

16. Wang, W. et al. Distinct cerebellar engrams in short-term and long-term motor learning. Proc. Natl. Acad. Sci. U. S. A. 111, E188–93 (2014).

17. Hippenmeyer, S., Johnson, R. L. & Luo, L. Mosaic Analysis with Double Markers Reveals Cell-Type-Specific Paternal Growth Dominance. Cell Rep. 3, 960–967 (2013).

18. Walker, J. H. & Agoston, D. V. The synaptic vesicle and the cytoskeleton. Biochem. J 247, 249–258 (1987).

19. Hirokawa, N., Niwa, S. & Tanaka, Y. Molecular Motors in Neurons: Transport Mechanisms and Roles in Brain Function, Development, and Disease. Neuron 68, 610–638 (2010).

20. Niranjan, P. S. et al. Thermodynamic regulation of actin polymerization. J. Chem. Phys. 114, 10573–10576 (2001).

21. Hanley, J. G. Actin-dependent mechanisms in AMPA receptor trafficking. Front. Cell. Neurosci. 8, 381 (2014).

22. Landis, D. M. & Reese, T. S. Differences in membrane structure between excitatory and inhibitory synapses in the cerebellar cortex. J. Comp. Neurol. 155, 93–125 (1974).

23. Harris, K. M. & Landis, D. M. Membrane structure at synaptic junctions in area CA1 of the rat hippocampus. Neuroscience 19, 857–872 (1986).

24. Masugi-Tokita, M. et al. Number and Density of AMPA Receptors in Individual Synapses in the Rat Cerebellum as Revealed by SDS-Digested Freeze-Fracture Replica Labeling. Journal of Neuroscience 27, 2135–2144 (2007).

25. Miyazaki, T., Fukaya, M., Shimizu, H. & Watanabe, M. Subtype switching of vesicular glutamate transporters at parallel fibre--Purkinje cell synapses in developing mouse cerebellum. Eur. J. Neurosci. 17, 2563–2572 (2003). 2+

26. Miki, T. et al. Numbers of presynaptic Ca^2+^ channel clusters match those of functionally defined vesicular docking sites in single central synapses. Proceedings of the National Academy of Sciences 201704470 (2017).

27. Kakegawa, W. et al. Optogenetic Control of Synaptic AMPA Receptor Endocytosis Reveals Roles of LTD in Motor Learning. Neuron 99, 985–998.e6 (2018).

28. Schonewille, M. et al. Zonal organization of the mouse flocculus: physiology, input, and output. J. Comp. Neurol. 497, 670–682 (2006).

29. Inoshita, T. & Hirano, T. Occurrence of long-term depression in the cerebellar flocculus during adaptation of optokinetic response. Elife 7, (2018).

30. Siksou, L., Triller, A. & Marty, S. An emerging view of presynaptic structure from electron microscopic studies. J. Neurochem. 108, 1336–1342 (2009).

31. Schrod, N. et al. Pleomorphic linkers as ubiquitous structural organizers of vesicles in axons. PLoS One 13, e0197886 (2018).

32. Schikorski, T. & Stevens, C. F. Morphological correlates of functionally defined synaptic vesicle populations. Nat. Neurosci. 4, 391–395 (2001).

33. Kaeser, P. S. & Regehr, W. G. The readily releasable pool of synaptic vesicles. Current Opinion in Neurobiology 43, 63–70 (2017).

34. Cingolani, L. A. & Goda, Y. Actin in action: The interplay between the actin cytoskeleton and synaptic efficacy. Nat. Rev. Neurosci. 9, 344–356 (2008).

35. Bissen, D., Foss, F. & Acker-Palmer, A. AMPA receptors and their minions: auxiliary proteins in AMPA receptor trafficking. Cell. Mol. Life Sci. 76, 2133–2169 (2019).

36. Fiala, J. C. & Harris, K. M. Cylindrical diameters method for calibrating section thickness in serial electron microscopy. J. Microsc. 202, 468–472 (2001).

37. Nusser, Z. et al. Cell type and pathway dependence of synaptic AMPA receptor number and variability in the hippocampus. Neuron 21, 545–559 (1998).

38. Holderith, N. et al. Release probability of hippocampal glutamatergic terminals scales with the size of the active zone. Nat. Neurosci. 15, 988–997 (2012).

39. Nakamura, Y. et al. Nanoscale Distribution of Presynaptic Ca2+Channels and Its Impact on Vesicular Release during Development. Neuron 85, 145–159 (2015).

40. Foster, K. A. & Regehr, W. G. Variance-Mean Analysis in the Presence of a Rapid Antagonist Indicates Vesicle Depletion Underlies Depression at the Climbing Fiber Synapse. Neuron 43, 119–131 (2004).

41. Guzman, S. J., Schlögl, A. & Schmidt-Hieber, C. Stimfit: quantifying electrophysiological data with Python. Front. Neuroinform. 8, 16 (2014).

42. Jonas, P., Major, G. & Sakmann, B. Quantal components of unitary EPSCs at the mossy fibre synapse on CA3 pyramidal cells of rat hippocampus. J. Physiol. 472, 615–663 (1993).

