## Supplementary information for "Advantages of acute brain slices prepared at physiological temperature in characterization of synaptic functions"

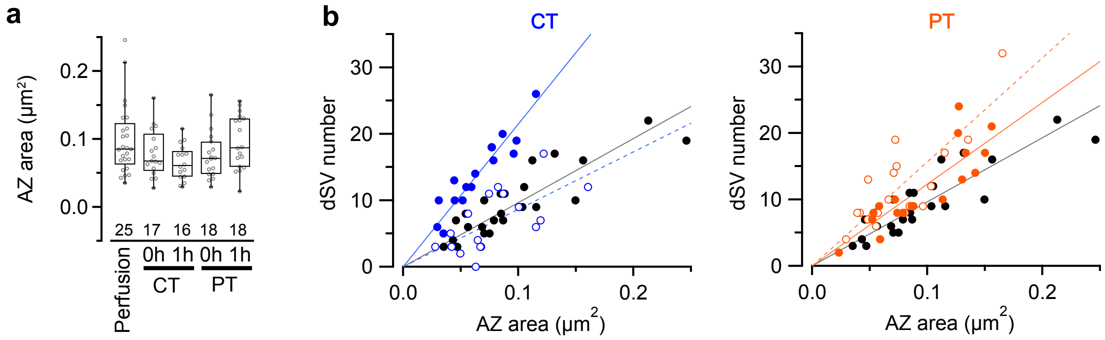


**Supplementary Figure 1**

Correlation between AZ area and docked SV numbers

**a,** Summary of the AZ area in PF boutons. Numerals in plot indicate the numbers of analyzed boutons for each group.

**b,** Correlation between AZ area and dSV number in cold-cut (left) or warm-cut (right) slices. Black circles indicate the values in perfusion-fixed tissues. Open and closed circles indicate before and after 1-h recovery time, respectively. Lines show the regression lines of each group (blue: CT, red: PT, dotted: 0 h, solid: 1 h).

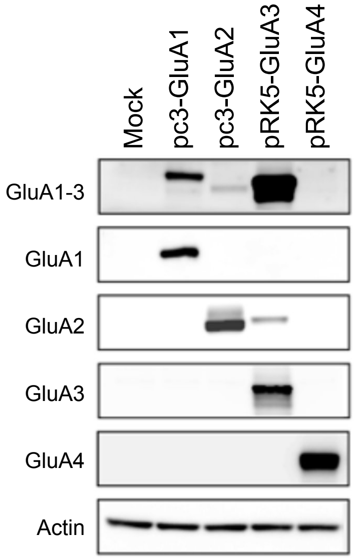


**Supplementary Figure 2**

Specificity of anti-GluA1-3 antibody validated by Western blot analysis

COS-7 cells were transfected with expression vector for AMPA receptor subunits and analyzed by immunoblotting with anti-GluA1-3, anti-GluA1, anti-GluA2, anti-GluA3, anti-GluA4, or anti-actin antibody. The anti-GluA1-3 antibody reacted with GluA1, GluA2 and GluA3, but not with GluA4.

**Table S1.** Electrophysiological properties of PF-PC synapses in mouse cerebellar slices prepared at CT and PT

| | CT | | | PT | | | |
|---|---|---|---|---|---|---|---|
| | n¶ | median | IQR | n¶ | median | IQR | *P-value*† |
| **Evoked EPSC kinetics** | | | | | | | |
| amplitude (4-V stim, pA) | 9 | 372.8 | 190.3 - 418.2 | 9 | 491.8 | 306.4 - 619.5 | *0.344* |
| 10-90% rise time (ms) | 9 | 2.4 | 1.7 - 2.6 | 9 | 2.1 | 1.9 - 2.4 | *0.486* |
| Decay time constant (ms) | 9 | 12.2 | 10.9 - 13.7 | 9 | 12.3 | 12.1 - 12.7 | *0.822* |
| **Variance-mean analysis** | | | | | | | |
| Release probability | 8 | 0.10 | 0.07 - 0.15 | 8 | 0.20 | 0.16 - 0.23 | *0.011** |
| quantal size (pA) | 6 | 1.10 | 0.83 - 1.37 | 8 | 1.06 | 0.83 - 1.26 | *0.963* |

†Welch's *t*-test

¶Number of cells

**P < 0.05*

**Table S2.** Dendritic spine density of Purkinje cells in perfusion-fixed and acute cerebellar slice tissues.

| | n[¶] | median | IQR | *P-value*[†] | | | | |
|---|---|---|---|---|---|---|---|---|
| | | | | *vs Perfusion* | *vs CT/0h* | *vs CT/1h* | *vs PT/0h* | *vs PT/1h* |
| **Spine density ($\mu m^{-1}$)** | | | | | | | | |
| Perfusion | 12 | 8.02 | 7.50 - 8.54 | - | | | | |
| CT/0h | 12 | 4.64 | 3.67 - 5.81 | *<0.001*** | - | | | |
| CT/1h | 11 | 9.30 | 7.85 - 9.79 | *0.651* | *<0.001*** | - | | |
| PT/0h | 10 | 7.11 | 6.49 - 7.90 | *0.718* | *0.043** | *0.097* | - | |
| PT/1h | 14 | 7.98 | 6.62 - 9.69 | *1* | *<0.001*** | *0.722* | *0.592* | - |

[†]One-way ANOVA with *post-hoc* Tukey-Kramer test

[¶]Number of dendrites

**P < 0.05, **P < 0.01*

**Table S3.** Synaptic vesicle distribution at PF boutons in perfusion-fixed and acute cerebellar slice tissues.

| | n[¶] | median | IQR | *P-value*[†] *vs Perfusion* | *vs CT/0h* | *vs CT/1h* | *vs PT/0h* | *vs PT/1h* |
|---|---|---|---|---|---|---|---|---|
| **AZ area (μm²)** | | | | | | | | |
| Perfusion | 25 | 0.085 | 0.069 - 0.131 | - | | | | |
| CT/0h | 17 | 0.068 | 0.056 - 0.100 | *1* | - | | | |
| CT/1h | 16 | 0.061 | 0.045 - 0.080 | *0.260* | *1* | - | | |
| PT/0h | 18 | 0.071 | 0.050 - 0.093 | *1* | *1* | *1* | - | |
| PT/1h | 18 | 0.087 | 0.062 - 0.133 | *1* | *1* | *0.280* | *1* | - |
| **Total SV number** | | | | | | | | |
| Perfusion | 16 | 137.0 | 73.0 - 177.8 | - | | | | |
| CT/0h | 12 | 160.0 | 110.8 - 177.3 | *1* | - | | | |
| CT/1h | 13 | 229.0 | 196.0 - 236.0 | *0.065* | *0.106* | - | | |
| PT/0h | 14 | 139.5 | 93.5 - 224.3 | *1* | *1* | *0.863* | - | |
| PT/1h | 17 | 166.0 | 149.0 - 272.0 | *0.838* | *1* | *1* | *1* | - |
| **dSV number** | | | | | | | | |
| Perfusion | 25 | 9.0 | 6.0 - 16.0 | - | | | | |
| CT/0h | 17 | 6.0 | 3.0 - 11.0 | *0.74* | - | | | |
| CT/1h | 16 | 13.5 | 10.0 - 17.3 | *0.15* | *0.006*** | - | | |
| PT/0h | 18 | 9.5 | 8.0 - 14.8 | *1* | *0.094* | *1* | - | |
| PT/1h | 18 | 9.5 | 8.0 - 17.0 | *1* | *0.205* | *1* | *1* | - |
| **dSV density (μm⁻²)** | | | | | | | | |
| Perfusion | 25 | 94.0 | 80.6 - 121.9 | - | | | | |
| CT/0h | 17 | 74.8 | 52.0 - 126.6 | *1* | - | | | |
| CT/1h | 16 | 221.0 | 200.1 - 232.1 | *<0.001*** | *<0.001*** | - | | |
| PT/0h | 18 | 146.1 | 121.0 - 196.6 | *0.0012*** | *0.005*** | *0.025** | - | |
| PT/1h | 18 | 110.9 | 100.2 - 138.0 | *0.631* | *0.361* | *<0.001*** | *0.109* | - |

[†]Kruskal-Wallis *H* test with *post-hoc* Mann-Whitney *U*-test with Bonferroni correction

[¶]Number of boutons

**P < 0.05, **P < 0.01*

**Table S4.** Immunogold particle density on PSD/AZs of PF-PC synapses in perfusion-fixed and acute cerebellar slice tissues.

| | n[¶] | median | IQR | *P-value*[†] | | | | |
|---|---|---|---|---|---|---|---|---|
| | | | | *vs Perfusion* | *vs CT/0h* | *vs CT/1h* | *vs PT/0h* | *vs PT/1h* |
| **GluA1-3 (μm$^{-2}$)** | | | | | | | | |
| Perfusion | 5 | 373.4 | 344.1 - 422.6 | - | | | | |
| CT/0h | 8 | 252.3 | 170.9 - 326.2 | *0.196* | - | | | |
| CT/1h | 8 | 213.0 | 161.6 - 257.0 | *0.048** | *0.933* | - | | |
| PT/0h | 7 | 338.7 | 241.1 - 419.9 | *0.926* | *0.545* | *0.173* | - | |
| PT/1h | 7 | 268.1 | 231.9 - 426.6 | *0.739* | *0.818* | *0.374* | *0.991* | - |
| **GluRD2 (μm$^{-2}$)** | | | | | | | | |
| Perfusion | 8 | 902.2 | 852.9 - 1072.5 | - | | | | |
| CT/0h | 6 | 662.4 | 605.7 - 724.7 | *0.104* | - | | | |
| CT/1h | 6 | 574.2 | 541.6 - 745.0 | *0.077* | *1* | - | | |
| PT/0h | 6 | 824.3 | 688.3 - 1000.5 | *0.874* | *0.562* | *0.478* | - | |
| PT/1h | 6 | 769.2 | 682.9 - 1033.5 | *0.718* | *0.735* | *0.653* | *0.998* | - |
| **RIM1/2 (μm$^{-2}$)** | | | | | | | | |
| Perfusion | 8 | 229.7 | 170.8 - 318.7 | - | | | | |
| CT/0h | 7 | 121.6 | 70.7 - 139.1 | *0.017** | - | | | |
| CT/1h | 7 | 219.5 | 182.3 - 235.9 | *0.88* | *0.15* | - | | |
| PT/0h | 5 | 172.7 | 167.4 - 237.0 | *0.599* | *0.517* | *0.976* | - | |
| PT/1h | 5 | 245.8 | 207.3 - 279.1 | *1* | *0.052* | *0.948* | *0.74* | - |
| **$Ca_V$2.1 (μm$^{-2}$)** | | | | | | | | |
| Perfusion | 8 | 219.3 | 176.5 - 241.6 | - | | | | |
| CT/0h | 7 | 133.8 | 88.5 - 143.7 | *0.005*** | - | | | |
| CT/1h | 7 | 137.9 | 101.4 - 166.7 | *0.026** | *0.962* | - | | |
| PT/0h | 5 | 160.2 | 151.4 - 170.0 | *0.298* | *0.569* | *0.895* | - | |
| PT/1h | 5 | 152.6 | 136.3 - 162.0 | *0.563* | *0.309* | *0.657* | *0.993* | - |

[†]One-way ANOVA with *post-hoc* Tukey-Kramer test

[¶]Number of replicas

**P < 0.05, **P < 0.01*

**Table S5.** Long-term depression on PF-PC synapses induced by HOKR training.

| | Control | | | HOKR-trained | | | |
|---|---|---|---|---|---|---|---|
| | n¶ | median | IQR | n¶ | median | IQR | *P-value*† |
| **mEPSC amplitude (pA)** | | | | | | | |
| CT | 14 | 23.34 | 19.96 - 25.52 | 17 | 20.70 | 18.29 - 25.59 | *0.154* |
| PT | 10 | 24.03 | 23.24 - 24.55 | 14 | 19.11 | 16.70 - 21.06 | *<0.001*** |
| | *P-value*† | *0.866* | | *P-value*† | *0.278* | | |
| **mEPSC frequency (Hz)** | | | | | | | |
| CT | 14 | 2.46 | 1.10 - 3.24 | 17 | 1.50 | 0.96 - 1.66 | *0.056* |
| PT | 10 | 0.29 | 0.26 - 0.38 | 14 | 0.10 | 0.07 - 0.51 | *0.626* |
| | *P-value*† | *<0.001*** | | *P-value*† | *0.005*** | | |

†Welch's *t*-test

¶Number of cells

$**P < 0.01$
